# genomeview - an extensible python-based genomics visualization engine

**DOI:** 10.1101/355636

**Authors:** Noah Spies, Justin Zook, Arend Sidow, Marc Salit

## Abstract

Visual inspection and analysis is integral to quality control, hypothesis generation, methods development and validation of genomic data. The richness and complexity of genomic data necessitates customized visualizations highlighting specific features of interest while hiding the often vast tide of irrelevant attributes. However, the majority of genome-visualization occurs either in general-purpose tools such as IGV (Robinson et al, 2011) or the UCSC Genome Browser (Kent et al, 2002) - which offer many options to adjust visualization parameters, but very little in the way of extensibility - or narrowly-focused tools aiming to solve a single visualization problem. Here, we present genomeview, a python-based visualization engine which is easy to extend and simple to integrate into existing analysis pipelines.

## Description

Genomeview includes built-in support for visualizing a number of standard genomics data types:

- Genomic features such as transcripts, exons and introns
- Sequencing data including reads, read-pairs, mismatches and insertions/deletions (indels)
- Quantitative data specified at the nucleotide or locus level

**Figure 1.**
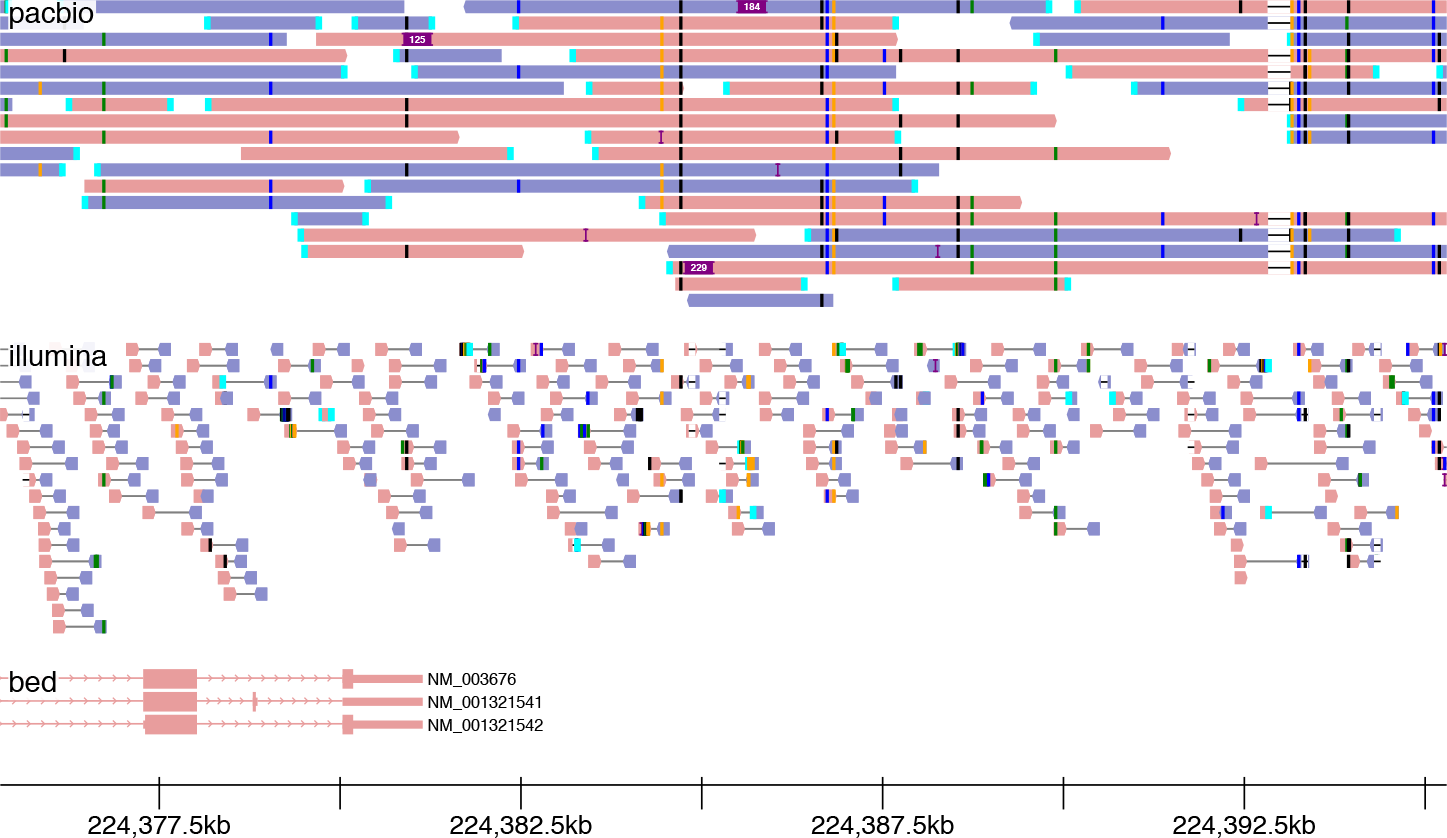
Genomeview visualization of PacBio (top), paired-end Illumina (middle) and a BED-format annotation track (bottom). Mismatches (here, mostly SNPs) are shown as short vertical bars. Insertions (here, mostly sequencing errors) are shown as purple bars with text indicating the length. A deletion is depicted by thin lines connecting PacBio sequences. Reads are colored blue or pink based on the mapped strand.

These features can be provided directly from python or in accompanying files in BAM/CRAM, BED and BigWig formats. In addition, genomeview includes several features intended to improve the visualization of noisy long-fragment sequencing data such as PacBio and Oxford Nanopore, most notably a quick-consensus mode that hides putative sequencing errors while retaining likely variants.

Genomic features and data are visualized relative to a common reference coordinate system, typically a reference genome. Notably, the python interface to genomeview facilitates visualizing novel nonreference genomic loci, for example displaying features and support relative to contigs from a genomic sequence assembly.

Genomeview natively outputs to the web graphics standard SVG file format, making it trivial to produce and view visualizations from the jupyter web-based interactive data analysis platform (Kluyver et al, 2016). In addition, genomeview can also output to PDF or PNG format, enabling the generation of visualizations as an automated step during analysis pipelines.

Genomeview takes advantage of the vibrant bioinformatics ecosystem in python. Of particular interest, because genomeview builds on pysam (itself built on htslib) and pyBigWig, most native data types can be streamed over the internet and thus only require downloading a small index file plus the data in the genomic region of interest.

Finally, genomeview is intended as a platform for customized genomic visualizations. The simple, object-oriented python implementation includes opportunities for python callbacks, enabling, for example, programmatically selecting and highlighting reads of interest; inheritance of the included track types in order to more deeply modify visual output; and creation of entirely new visualization tracks.

## Usage

An example genomeview visualization demonstrates the automatic filtering of erroneous mismatches and short indels from a PacBio dataset, emphasizing a small number of single nucleotide polymorphisms (SNPs) present in the sample (Figure 1). The code and data necessary to produce visualizations is available online from github and using the jupyter nbviewer at http://bit.ly/2Ipfs33 (also available as Supplementary Figure 1). Data is drawn from Genome in a Bottle sample HG002 (Zook et al, 2018).

That notebook demonstrates how substantial new functionality can be obtained with just a few additional lines of code. Up-to-date documentation can be viewed at http://genomeview.readthedocs.io/.

## Discussion

Genomeview has substantial utility for both high-throughput visualization of many loci and datasets directly from python, as well for customizable views of genomic data. As the underlying visualization engine for the latest version of svviz (Spies et al, 2015), genomeview is already being used widely in a production setting. In addition, we have found it to be invaluable in an exploratory research context, allowing us to easily and quickly create customized and focused views of highly error-prone long-fragment sequencing data and whole-genome sequence assembly results directly from within jupyter.

## Availability

Genomeview is available as a python package from PyPI (https://pypi.org/project/genomeview) and the source code is distributed under the MIT license (https://github.com/nspies/genomeview). Questions, feature requests, bug reports and pull requests can be submitted on the github issues page (https://github.com/nspies/genomeview/issues).

## Acknowledgements

Certain commercial equipment, instruments, or materials are identified to adequately specify experimental conditions or reported results. Such identification does not imply recommendation or endorsement by the National Institute of Standards and Technology, nor does it imply that the equipment, instruments, or materials identified are necessarily the best available for the purpose.

